# High frequency oscillations and the 1/f slope vary across a spectrum of depression severity

**DOI:** 10.1101/2025.07.27.667095

**Authors:** Juliet Hosler, James Coxon, Joshua Hendrikse

## Abstract

Depression is a highly prevalent mental disorder that impacts an individual’s functioning, societal productivity, and quality of life. It is associated with disrupted neural activity (e.g., balance of excitation-inhibition) across networks implicated in emotional processing, such as between prefrontal and limbic regions. High frequency oscillations measured with electroencephalography (i.e., beta and gamma rhythms) differ between healthy individuals and those with depression, and predict treatment outcomes following pharmacological intervention. However, to date research has focussed on binary comparisons between individuals with a clinical depression diagnosis relative to healthy control populations, providing limited insight into how these measures may shift as a function of illness severity. To establish the utility of EEG measures as potential biomarkers for depression, an improved understanding across the spectrum of symptom profiles is required. Here, we aimed to bridge this gap and investigate changes in high frequency beta and gamma oscillations and the 1/f slope in resting-state EEG across a spectrum of mild to severe depression symptom presentations. In line with expectations, we demonstrate graded alterations to gamma and beta power and the 1/f slope across a spectrum of depression severity. Our findings provide critical new insights into the neurophysiological signature of depression symptoms, and highlight the utility of EEG markers to inform future precision psychiatry approaches to more effectively assess and treat depression.

**Highlights:**

- High frequency brain oscillations are associated with depression severity
- Excitation-inhibition balance is altered in depression
- These neural markers could inform precision psychiatry approaches

## Introduction

Depression is a highly prevalent mental disorder characterised by low mood and/or anhedonia, which can have devastating impacts on daily functioning, social engagement, and occupational productivity (American Psychiatric Association, 2013; McTernan et al., 2013). Although depression symptoms are notoriously heterogeneous (Fried & Nesse, 2015), neuroimaging studies indicate they are commonly associated with disruptions across networks implicated in emotional processing (e.g., frontolimbic regions; for review, see: Zhang et al., 2018). Identification of neural markers that reflect communication within these neural circuits may assist in tailoring treatment approaches to individual clinical presentations.

EEG is an accessible way to measure neural communication. The EEG signal at a given electrode is comprised of both periodic, oscillatory activity, and aperiodic activity. The overall signal can be decomposed into the relative contribution of each frequency band in the form of a power spectrum. Different compositions of spectral power underpin neural communication between spatially distinct brain regions (Fries, 2005; Izhikevich, 1999, see Figure 1A (left)). For example, synchronisation of high frequency oscillations facilitates emotional expression recognition by integrating information from several spatially distinct brain regions (Symons et al., 2016). Individuals with depression have exhibited disruption in connectivity between brain regions involved in processes such as emotional regulation, emotional perception, and social cognition (Guo et al., 2016), likely related to aberrant oscillatory communication. In particular, high frequency oscillations have garnered interest as a possible marker of depression severity (Fitzgerald & Watson, 2018; Gilbert & Zarate, 2020).

**Figure 1.**
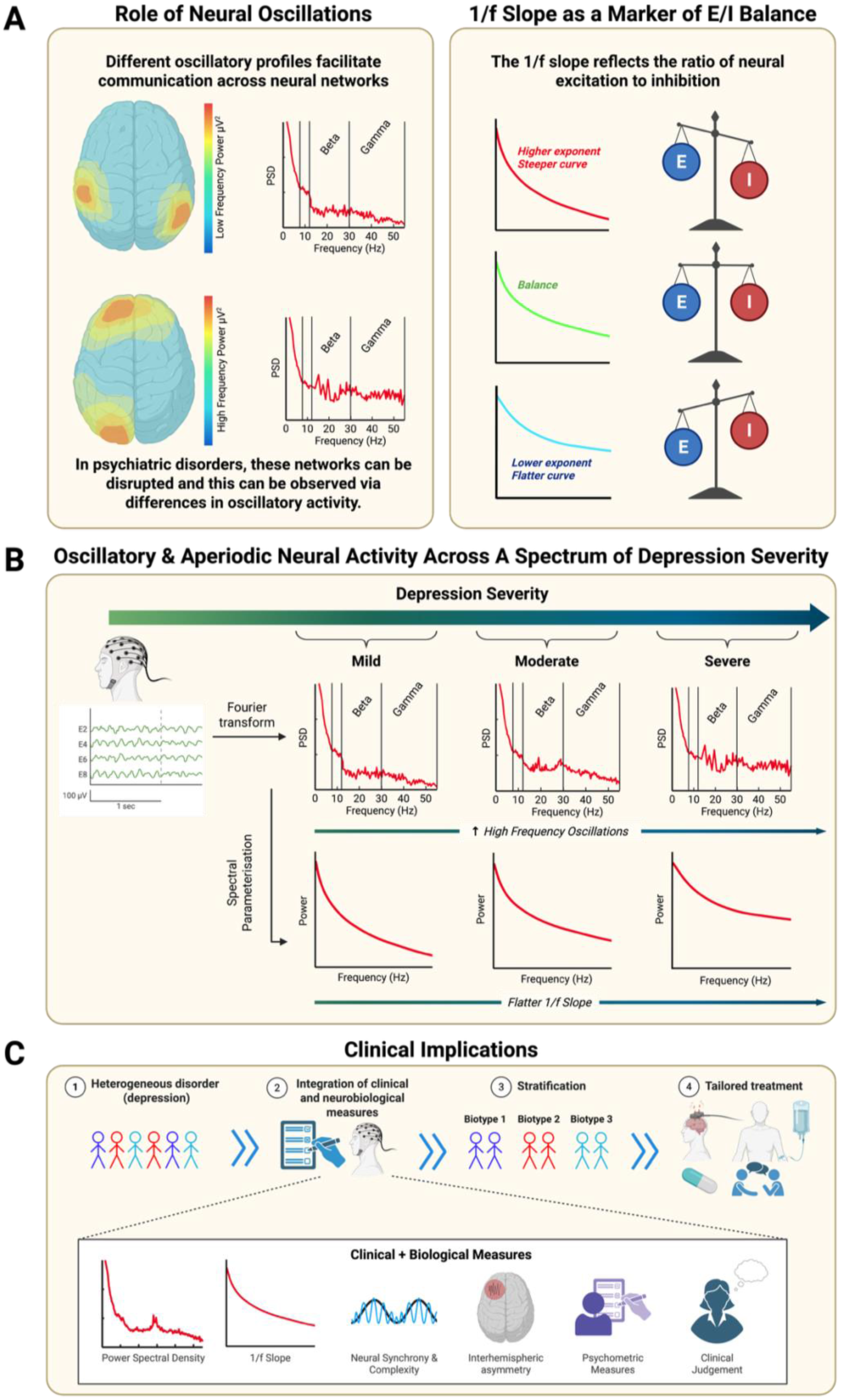
Theoretical framework for mapping EEG features to depression symptomology. A (left): Disruptions in neural circuits can be characterised via analysis of oscillatory activity measured by non-invasive EEG. Past studies have identified elevated high frequency (e.g., gamma) activity in individuals with depression. A (right): The 1/f slope provides an electrophysiological measure of excitation-inhibition balance, with steeper slopes reflecting greater inhibition and flatter slopes reflecting greater excitation. B: Schematic representing the current study’s methodology and hypotheses. EEG data is transformed into the frequency domain, providing a power spectrum outlining the contributions of relative frequencies. This data also undergoes “spectral parameterisation”, in which the aperiodic component of the signal (1/f slope) is extracted. High frequency oscillations and the 1/f slope are expected to be sensitive to increasing depression severity. C: The current study contributes to efforts to better tailor treatment to individuals by providing potential markers that, in conjunction with other established measures, can stratify individuals into subgroups based on their neural characteristics.

High frequency ‘gamma’ oscillations (30 Hz and greater) index the rhythmic firing of inhibitory interneurons (Sohal et al., 2009), which occur as part of a negative feedback loop in the communication between excitatory and inhibitory neurons (Sohal, 2016). Thus, gamma oscillations are thought to reflect a region being “active” (Merker, 2013), and have also been used as a measure of the ratio of excitation to inhibition across the cortex (Snijders et al., 2013). Gamma oscillations are involved in numerous cognitive and affective processes relevant to depression such as perceptual awareness, attention, memory, (Sedley & Cunningham, 2013), and emotional processing (Liu et al., 2014; Siegle et al., 2007).

Several studies have identified significant differences in resting-state gamma power when comparing depression and healthy control groups (Liu et al., 2022; Tekell et al., 2005; Zeng et al., 2024; for review, see Fitzgerald & Watson, 2018), and also demonstrated that gamma power differentiates individuals with depression from those with other psychiatric diagnoses, such as bipolar disorder (Liu et al., 2012; Liu et al., 2014). Further, gamma power is altered following antidepressant treatments, such as ketamine and repetitive transcranial magnetic stimulation (Noda et al., 2017; Nugent et al., 2019; Pathak et al., 2016). However, there is also evidence that oscillatory changes in depression are not specific to gamma. Differences in the beta band (∼14 Hz to 30 Hz) have also been observed between depression and healthy controls (Grin-Yatsenko et al., 2009, 2010; Knott et al., 2001), and following ketamine treatment (Anijärv et al., 2023). Beta power is thought to relate to working memory and attention, which are negatively impacted in depression (Li et al., 2017), and increased beta power is correlated with the number of depressive episodes and refractory depression (Grin-Yatsenko et al., 2009, 2010; Knott et al., 2001; Matousek, 1991; Nystrom et al., 1986).

Taken together, these findings suggest that high frequency oscillations (gamma and/or beta) could be a viable neural marker for identifying participants with depression. However, it remains unclear how these markers may map onto the spectrum of depression severities.

In addition to oscillatory patterns, the non-periodic 1/f slope of the EEG signal provides a useful measure of excitation-inhibition balance across the cortex (Donoghue et al., 2020) (Gao et al., 2017), see Figure 1A (right). The slope is extracted by applying a model to the frequency transformed EEG data and can be described by its two parameters, the exponent (curvature of the slope) and offset (y-intercept), where the slope reflects characteristic differences between excitatory and inhibitory current decay and the offset reflects broadband shifts in power (Donoghue et al., 2020, see Figure 1A (right)). Differences in E/I balance have been reported between individuals with a history of depression relative to healthy controls. However, measurements to date have been taken from either post-mortem tissue cell counts (Hu et al., 2023), or via magnetic resonance spectroscopy (Kantrowitz et al., 2021; Romeo et al., 2018; Schür et al., 2016) with the latter providing an overall estimate of neurometabolite concentration that is thought to predominantly index metabolic and extrasynaptic processes (Stagg et al., 2011). Thus, the 1/f slope may provide an alternative non-invasive way to index synaptic E/I balance, and examine corresponding changes in E/I balance across depression severities.

To our knowledge, only one study has examined the role of the 1/f slope in depression (Hacker et al., 2025). Using intracranial electrodes in five individuals with refractory depression, the study found initial evidence for a relationship between the 1/f slope and depression severity, with a flatter curve (greater excitation) indicating reduced depression severity. Intracranial electrodes are not feasible for use in the general depression population due to their invasive nature, and larger sample sizes may yield more reliable inference. Thus, incorporating non-invasive techniques to assess changes in E-I balance may offer novel insights into the neurophysiology associated with distinct clinical presentations (see Figure 1B).

An improved understanding of how neural transmission is altered as a function of depression severity may advance screening and assessment of depression and facilitate more effective intervention (see Figure 1C). To this end, the present study aimed to assess changes in high frequency oscillations (i.e., beta and gamma power) and E-I balance across a spectrum of depression severities (see Figure 1B) using a large open-source dataset (the Brainclinics Foundation (van Dijk et al., 2022)). We hypothesised that both beta and gamma power would differ across individuals experiencing mild, moderate, and severe depression symptoms. We also hypothesised that the 1/f slope, a marker of E-I balance, would significantly differ across depression severities. To assess the reproducibility and robustness of observed effects, we examined these same associations in an independent dataset.

## Methods

### Data Acquisition & Participants

This study utilised open-source data from the Brainclinics Foundation (https://www.synapse.org/TDBRAIN), titled ‘Two Decades Brainclinics Research Archive for Insights in Neurophysiology’ (van Dijk et al., 2022), referred to here as TDBRAIN (2022). The dataset contains resting-state data recorded from a 26 channel EEG montage and self-reported depression symptom data from 1274 participants with mental health diagnoses collected between 2001 to 2021 (van Dijk et al., 2022) (age range 6-83 years, *M* = 37.48 ±19.1 SD). BDI scores and eyes-open resting-state recordings (two minutes duration) were obtained from N =181 for analysis in the current study. We restricted our analyses to the 181 participants for whom Beck Depression Inventory II (BDI) scores were recorded.

### The Beck Depression Inventory II

The Beck Depression Inventory II (BDI-II) is a well-validated 21-item self-report measure of depression symptoms in adults and adolescents (Beck et al., 1996; Osman et al., 2004). For each item, respondents are instructed to select the option that best describes how they have been feeling during the past two weeks. Each item is scored from 0 to 3 (e.g., 0 = I do not feel sad; 1 = I feel sad much of the time; 2 = I am sad all the time; 3 = I am so sad or unhappy that I can’t stand it). Final scores range from 0 to 63, with 0-13 classed as “minimal”, 14-19 as “mild”, 20-28 as “moderate”, and 29 and above as “severe” (Beck et al., 1996). The scale is regarded as a valid and reliable measure of depression symptoms (Dozois et al., 1998; Sprinkle et al., 2002), with an internal consistency of 0.9 and test-retest reliability from 0.73 to 0.96 (Wang & Gorenstein, 2013).

### Data Analysis

#### EEG Preprocessing

All EEG files were pre-processed using the Reduction of Electroencephalographic Artifacts (RELAX) automated pre-processing pipeline utilising EEGLAB functions within MATLAB (Bailey et al., 2023; Delorme & Makeig, 2004; The MathWorks Inc., 2023). Cerebellar, ocular, and mastoid electrodes were excluded and line noise was removed using ZaplinePlus (de Cheveigné, 2020; Klug & Kloosterman, 2022). The data was downsampled to 500 Hz, and bandpass filtered at (high-cutoff 0.5 Hz,120 Hz low-cutoff) with the pop_eegfiltnew function (Delorme & Makeig, 2004). Multi-Weiner Filtering (MWF) and Independent Component Analysis (ICA) (Picard algorithm) were employed to remove muscle and eye movement artifacts, with subsequent data interpolation. Default parameters were used for all other RELAX pre-processing steps. N = 8 were excluded on the basis of excessive noise remaining in resting-state EEG data following pre-processing.

#### Spectral Power Extraction

Pre-processed EEG files were imported into Brainstorm (2023), and the Welch Power spectrum density function was utilised to transform the data from the time domain to the frequency domain (window length = 4s, overlap ratio = 50%, units = physical μV^2^/Hz). Frequency bands were specified as theta: 4-7 Hz, alpha: 8-12 Hz, low beta: 13-20 Hz, high beta: 21-29 Hz, low gamma: 30-45 Hz, and high gamma: 55-90 Hz (avoiding 50 Hz line noise).

#### Fitting the 1/f Slope (Spectral Parameterisation)

1/f slopes were calculated from frequency transformed data using the ‘FOOOF: Fitting oscillations and 1/f’ package in Brainstorm (Donoghue et al., 2020). The frequency range for analysis was set at 1-40 Hz, with a gaussian peak model (peak width limits = 0.5-12 Hz, maximum number of peaks = 3, minimum peak height = 3 dB, proximity threshold = 2 standard deviations (SD) of peak model, fixed aperiodic model), yielding individual estimates of the aperiodic exponent and offset.

#### Cluster-based Permutation Statistical Analyses

For group-level analyses, outliers (+3SD) were identified independently per dependent measure (beta and gamma frequency bands, 1/f exponent and offset values). Descriptive statistics after outlier removal can be found in Table 1.

To identify differences in spectral power and the 1/f slope as a function of depression symptomology, cluster-based permutation analysis were conducted using the Monte Carlo method in the MATLAB-based toolbox FieldTrip (Oostenveld et al., 2011). Cluster-based Spearman correlations (two-tailed) were calculated between BDI-II scores and EEG outcomes of interest to assess associations between EEG measures (gamma and beta power, and 1/f estimates) and depression symptoms (cluster statistic = “maxsum”, number of randomisations = 10000, alpha threshold = 0.05). Independent-samples t-tests were also conducted to identify differences between diagnostic categories (e.g., mild vs severe symptom presentations).

Inspection of the TDBRAIN (2022) dataset indicated ‘mild’ (n = 22), ‘moderate’ (n = 52), and ‘severe’ (n = 87) symptom presentations. Therefore, separate cluster-based independent t-tests were performed between these groups (i.e., mild vs moderate, mild vs severe, and moderate vs severe).

### Dataset 2

While the TDBRAIN (2022) dataset encompassed a variety of scores and a large number of participants, it has some limitations. 52 participants were listed as having taken medication between 0 and 15 hours prior to data collection, and EEG data was collected using a relatively low-density 26 electrode array. To test robustness of our findings and account for these limitations, we sourced a second dataset with a carefully characterised sample and a higher density electrode array. This dataset (herein referred to as “Cavanagh (2021)”) was sourced from the OpenNeuro platform and contains resting-state data recorded from a 60 channel 10-20 EEG montage and self-reported depression symptom data from 122 young adults aged 18-24 (M = 18.57 ±1.19 SD). Participants had no history of head trauma or seizures and no current psychoactive medication use. “Control” participants had stable low BDI scores (remaining <7 for 2-14 weeks), no self-reported history of Major Depressive Disorder, and no self-reported symptoms associated with mental illness (previously referred to as Axis I disorders). Participants in the “depression” group had stable high BDI scores (remaining ≥ 13 for 2-14 weeks), and many were administered the Structured Clinical Interview for DSM-5, revealing almost all primary diagnoses were depression or anxiety-related.

Each resting-state EEG recording was six minutes in duration, with alternating epochs of 1-minute of rest with eyes open, and one minute with eyes closed. A custom MATLAB (2023b) script was used to exclude data during the eyes-closed condition to ensure consistency between datasets. Data from 6 participants were excluded on the basis of missing resting-state data (n = 5) or BDI-II scores (n = 1), leaving a total of N=116.

Methodology for pre-processing and data analysis for this dataset was largely the same as for the TDBRAIN (2022) dataset, with the following exceptions:

- N = 3 participants were excluded due to excessive noise remaining in resting-state EEG data following pre-processing.
- In spectral power extraction, low gamma power was defined as 30-55 Hz and high gamma as 65-90 Hz in order to avoid 60 Hz line noise for this North American dataset.
- The sampling strategy for this dataset resulted in the presence of a ‘healthy’ control group (BDI mean = 1.78, SD = 1.70) and another group with moderate-severe depression symptoms (BDI mean = 22.44, SD = 4.70). To accommodate the sampling differences between datasets, cluster-based permutation independent t-tests were performed between participants with scores between 0 and 13 (‘minimal’ category, n

= 73), and participants with scores 20 and above (‘moderate’ and ‘severe’, n = 41). Correlations were performed across all participants, that is, with the inclusion of participants in the ‘mild’ category, n = 13.

## Results

### Dataset 1 – TDBRAIN

#### Demographics

N = 173 were included in the final analysis. Participants had a mean age of 40.36 +-13.59, range 14-78 years. There were no significant differences in age (M = 40, SD = 13-14) or biological sex (47-50% female) between sub-samples (i.e., between frequency bands). Control analysis indicated that there was no significant relationship between age and BDI scores (rho = -.05, *p* = .53).

#### Associations between gamma power and depression symptomology

Topographical representations of gamma power as a function of depression symptom severity is represented in Figure 2A-B. Increased gamma power was observed in individuals experiencing severe symptoms, relative to individuals with moderate symptoms across a central-posterior electrode cluster (low gamma: t = −2.74, p = .002, Figure 2C; high gamma: t = −2.64, p = .003, Figure 2D). Furthermore, a correlation analysis across the entire sample revealed that low gamma power was positively correlated with BDI scores (Figure 2E), particularly across a left anterior-central cluster (rho = .20, p = .01) and right posterior-central cluster (rho = .19, p = .04).

**Figure 2.**
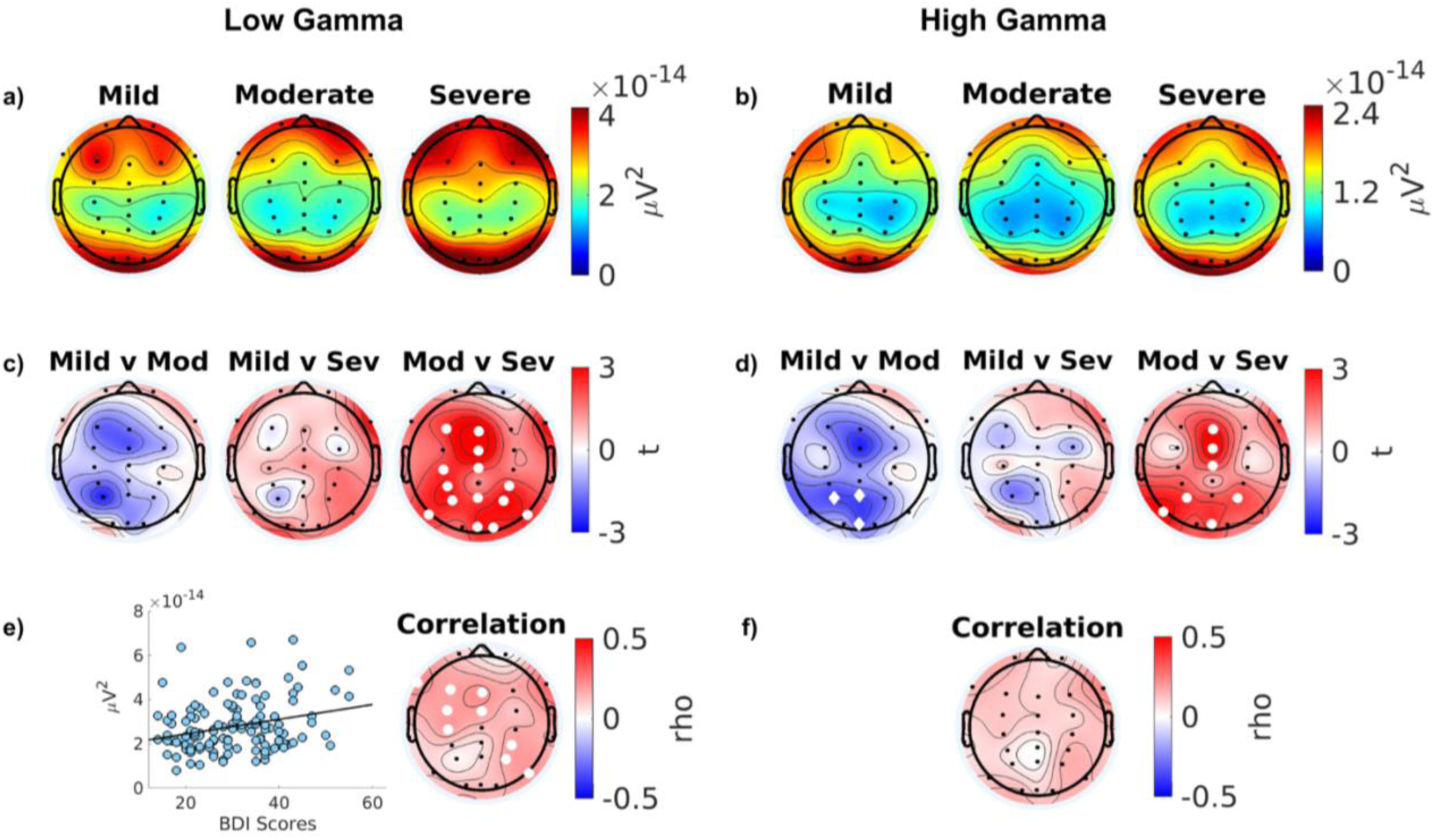
Topographical representation of gamma power across depression symptoms. Top row: low (a) and high (b) gamma power in mild, moderate, and severe BDI categories. Second row: t-statistics for comparisons of low (c) and high (d) gamma power between mild and moderate scorers, mild and severe scorers, and moderate and severe scorers. Third row: Cluster-based permutation Spearman correlations for low (e) and high (f) gamma power. Scatterplot depicts the correlation between BDI scores and average power across significant electrodes. For both t-test and correlation topography maps, red denotes an increase in power with greater depression severity, while blue denotes a decrease in power with greater depression severity. White circles: *p<.05, corrected for multiple comparisons. White diamonds: p<.05, uncorrected*.

#### Associations between beta power and depression symptomology

There was some evidence of increased beta power (both high and low) in individuals experiencing severe depression symptoms, relative to those with moderate symptoms (see Figure 3A&B). This difference was most pronounced in a central-anterior electrode cluster for high beta power (t = −2.22, p <.05 uncorrected) but did not remain statistically significant after adjusting for multiple comparisons (p = .07, Figure 3D). No other significant differences or associations were observed (Figure 3E&F).

**Figure 3.**
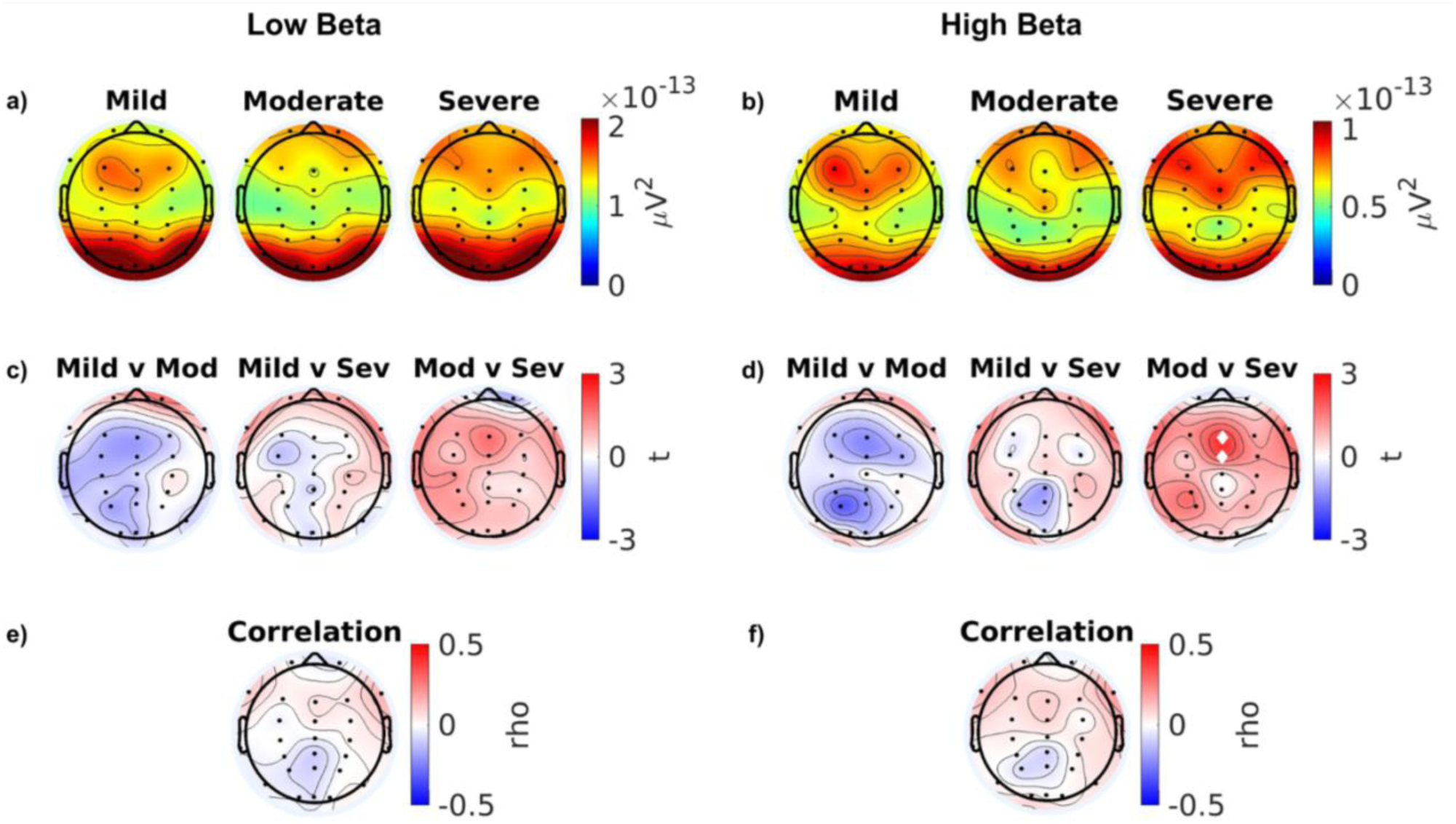
Topographical representation of beta power across depression symptoms. Topographical representation of beta power across depression symptoms. Top row: low (a) and high (b) beta power in mild, moderate, and severe BDI categories. Second row: t-statistics for comparisons of low (c) and high (d) beta power between mild and moderate scorers, mild and severe scorers, and moderate and severe scorers. Third row: Cluster-based permutation Spearman correlations for low (e) and high (f) beta power. Scatterplot depicts the correlation between BDI scores and average power across significant electrodes. For both t-test and correlation topography maps, red denotes an increase in power with greater depression severity, while blue denotes a decrease in power with greater depression severity. White circles: *p<.05, corrected for multiple comparisons. White diamonds: p<.05, uncorrected*.

#### Changes in the 1/*f* slope across differing depression severities

Topological representations of the spectral parameterisation (1/f exponent and offset) values for each sub-group are shown in Figure 4A-B. For the 1/f exponent, no significant group differences were observed (Figure 4C). There was some evidence of a positive correlation between depression symptoms and exponent values across a frontocentral topography (rho = .17, p < 0.03 uncorrected), though these results did not remain significant after correcting for multiple comparisons (p = .09, Figure 4E). A greater 1/f offset was observed in individuals with severe relative to moderate depression symptoms, across a centroparietal topography (t = −2.57, p = .007, Figure 4D). A significant positive correlation was also observed between offset and depression symptoms (rho = .21, p = .004, Figure 4F). Interestingly, the topography for this cluster was qualitatively similar to low gamma. As such, a correlation analysis was performed between low gamma power and 1/f offset for each electrode. This revealed significant moderate positive correlations for each electrode, with rho values spanning from .29 to .65 (see Figure 5). No other significant differences were observed.

**Figure 4.**
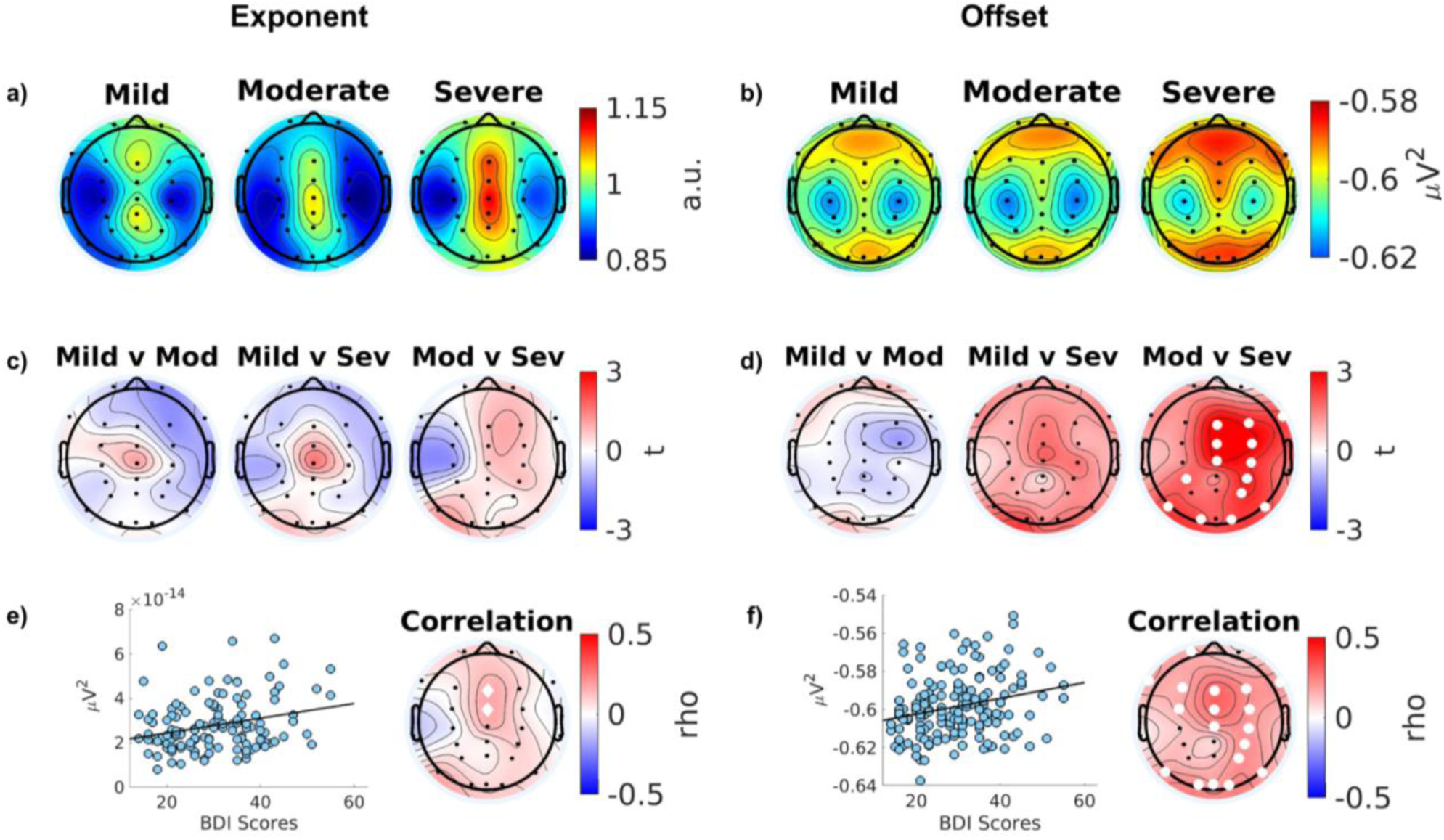
Topographical representation of 1/f parameters across depression symptoms. Top row: exponent (a) and offset (b) parameters in mild, moderate, and severe BDI categories. Second row: exponent (c) and offset (d) t-statistics for comparisons between mild and moderate scorers, mild and severe scorers, and moderate and severe scorers. Third row: Cluster-based permutation Spearman correlations for exponent (e) and offset (f) parameters. Scatterplot depicts the correlation between BDI scores and average parameter value across significant electrodes. For both t-test and correlation topography maps, red denotes an increase in parameter magnitude with greater depression severity, while blue denotes a decrease in parameter magnitude with greater depression severity. White circles: *p*<.05, corrected for multiple comparisons. White diamonds: *p*<.05, uncorrected.

**Figure 5.**
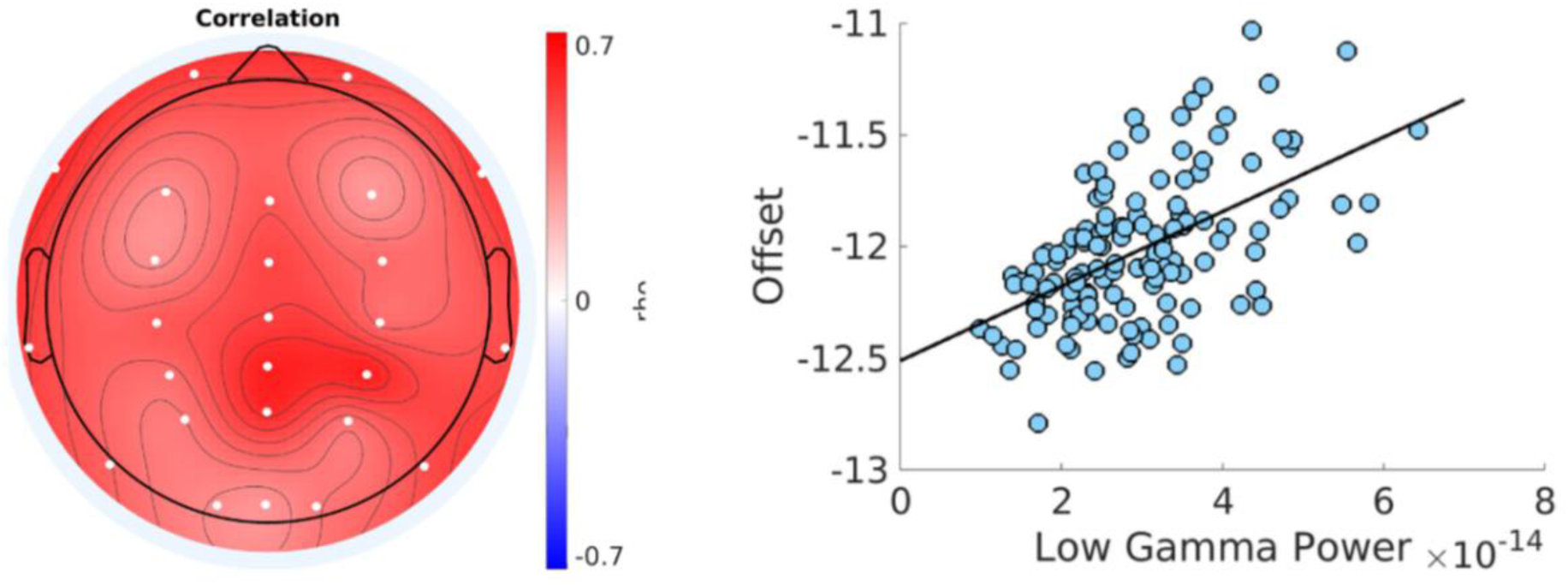
Topographical representation and scatter plot of correlation between low gamma power and 1/f offset. Left: Spearman’s rho values for the correlation between the two variables at each electrode. White circles: *p* < .017, Bonferroni-corrected. Right: Scatterplot depicting 1/f offset values against low gamma power values for each participant.

Overall, significant increases in gamma power and the offset parameter of the 1/f slope model were observed across depression severities, particularly between individuals with “moderate” and “severe” BDI scores. These effects had a distributed topography. We did not observe significant changes in the beta band, although there was a general trend towards increased beta power with increased depression severity. To assess the reproducibility of these findings, the analyses were also conducted on the Cavanagh (2021) dataset.

### Dataset 2 - Cavanagh

N = 113 were included in the final analysis. Participants had a mean age of 18.90 +-1.22, range 18-24 years. There were no significant differences in age (M = 19, SD = 1) or biological sex (38-53% female) between sub-samples (i.e., between frequency bands). Control analysis indicated that there was no significant relationship between age and BDI scores (rho = -.11, *p* = .25).

#### Associations between gamma power and depression symptomology

No significant differences in gamma power were observed between individuals with depression and controls for the Cavanagh dataset. There was evidence of a positive correlation between gamma power and BDI score in a left frontocentral and right posterior cluster, but this was a weak relationship (rho = .30, p < .02 uncorrected) that did not survive correction for multiple comparisons (p = .07), see Figure 6D).

**Figure 6.**
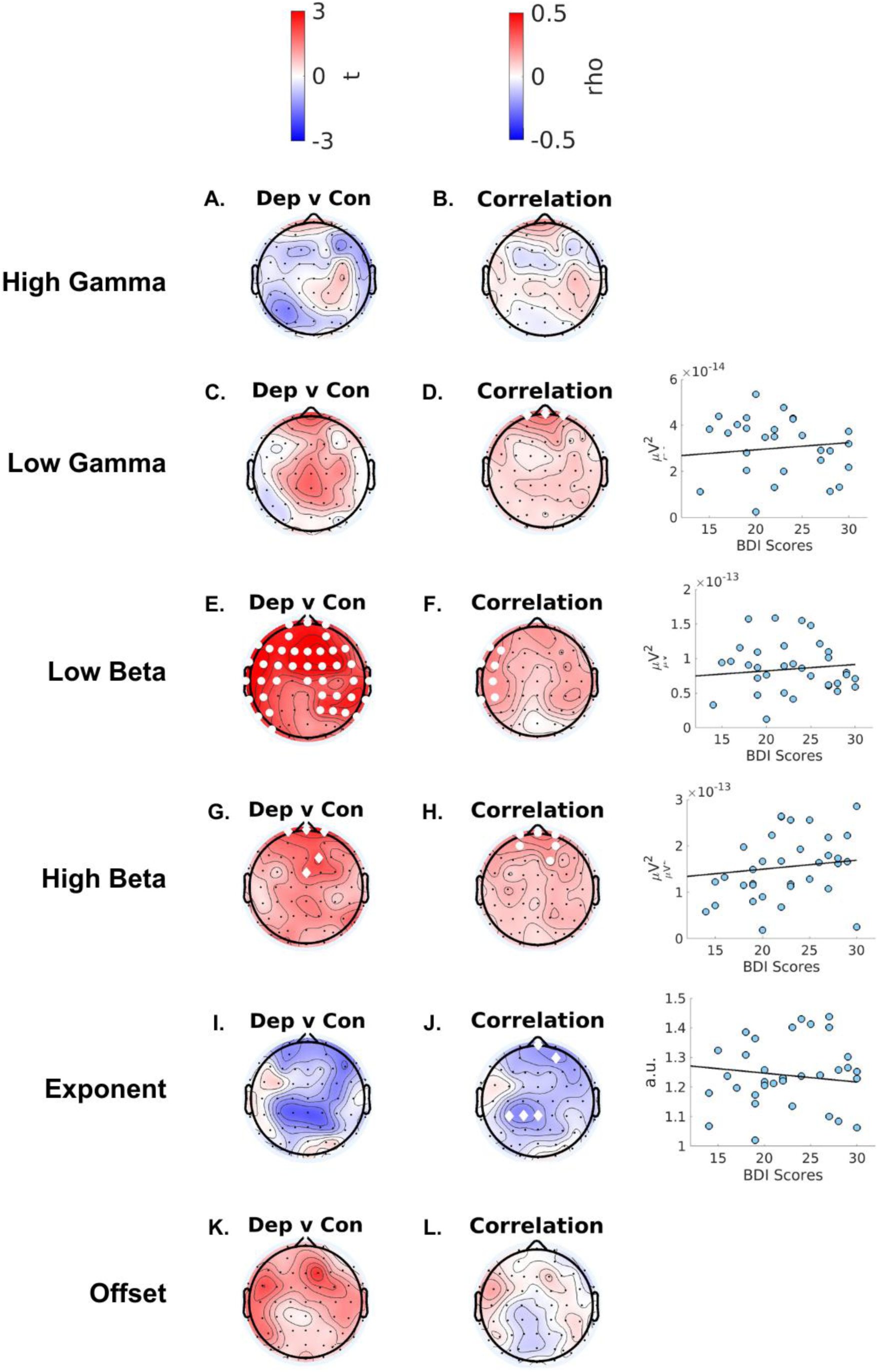
Topographical representation of comparisons and correlations of spectral power and 1/f slope parameters across depression symptoms. Rows, from top to bottom: t-statistics for comparisons between control group and depression group for spectral power and 1/f parameters, cluster-based permutation Spearman correlations for spectral power and 1/f parameters, and scatterplots depicting the correlation between BDI scores and average spectral power across significant electrodes. For both t-test and correlation topography maps, red denotes an increase in parameter magnitude with greater depression severity, while blue denotes a decrease in parameter magnitude with greater depression severity. White circles: *p* < .05, corrected for multiple comparisons. White diamonds: *p* < .05, uncorrected.

#### Associations between beta power and depression symptomology

Significantly higher low beta power was observed in individuals with depression relative to controls, across anterior, central, and temporal regions (t = −2.64, p = .005; see Figure 6E). Positive correlations between beta power and BDI scores were also observed. In the low beta band, after adjusting for multiple comparisons, this effect was most pronounced across a left temporal cluster (rho = .24, p = .047 corrected; see Figure 6F). In the high beta band, this effect was most prominent across an anterior cluster (rho = .24, p = .045 corrected; see Figure 6G).

#### Changes in the 1/*f* slope across differing depression severities

For the 1/f exponent, no significant group differences were observed (Figure 6I and 6J). There was some evidence of a negative correlation between depression symptoms and exponent values across an anterior (rho = -.22, p < .02, uncorrected) and central-posterior (rho = -.22, p < .03., uncorrected) topography, though these results did not remain significant after correcting for multiple comparisons (anterior: p = .10, central-posterior: p = .13). No significant group difference or correlation was observed for the 1/f offset (Figure 6 K&L)..

## Discussion

In this study, we demonstrate changes in high frequency oscillatory activity across a spectrum of depression symptom severities. We show modulation of gamma and beta power as a function of depression severity, and alterations in excitation-inhibition balance. We assessed the reproducibility of these effects using an independent dataset, and observed evidence of altered high frequency oscillations, albeit with a smaller effect size. To our knowledge this study is the first to demonstrate graded changes in oscillatory power across depression symptoms. We discuss the potential utility of these EEG features as possible markers of depression symptomology to aid in improved assessment and treatment.

### Gamma power is altered across depression severities

In line with our expectations, we observed altered gamma power across depression severities. Specifically, we demonstrate a general pattern of increased gamma power across increasing severity. Our findings align with previous work reporting differences in gamma power between individuals with depression and healthy controls when assessed at rest (Strelets et al., 2007; Tatti et al., 2024; Zeng et al., 2024), and during presentation of emotional stimuli (Lee et al., 2010; Liu et al., 2012; Liu et al., 2014; for review, see Fitzgerald & Watson, 2018). Extending on these findings, our results indicate that differences in high frequency oscillations are not only apparent between healthy individuals and those with a depression diagnosis, but show a relationship with depression symptom severity, with the most pronounced differences for moderate to severe symptom presentations.

Interestingly, we note certain inconsistencies between our findings of increasing gamma power and previous studies with regard to the direction of observed effects. For instance, Liu and colleagues reported decreased gamma power in drug-naïve individuals with first-episode depression (Liu et al. (2022). Chronicity (and by extension, history of medication/treatment) may influence the relationship between gamma activity and depression. Indeed, certain antidepressant treatments (such as rTMS and SSRIs) have been associated with increased gamma power (Noda et al., 2017; Pathak et al., 2016), and in certain cases change in gamma power has been associated with symptom remission (Pathak et al., 2016). Notably, we observed some differences in the strength of associations between datasets in the current study, which may also plausibly reflect differences in medication usage between samples. Overall, we show evidence that gamma power increases with increasing depression severity. Future work may explore the role of chronicity and treatment history in mediating the direction and magnitude of these associations.

### Greater depression severity is associated with greater beta power

We also observed evidence of increased beta power as a function of depression severity. Specifically, analysis of the primary dataset revealed preliminary evidence of increased high beta power in individuals with severe depression symptoms, relative to those with moderate symptom presentations. This finding was corroborated by analysis in an independent dataset, which similarly showed elevated beta power across a widespread topography in individuals with depression relative to healthy controls. Broadly, our findings align with several previous studies reporting altered beta power in individuals with depression across anterior (Knott et al., 2001; Suzuki et al., 1996), parieto-occipital (Grin-Yatsenko et al., 2009), and distributed (Grin-Yatsenko et al., 2010) topographies. Interestingly, in line with the current findings indicating a relationship between symptom severity and beta power, previous studies have also identified links between beta modulation and chronicity (e.g., relapse and number of depressive episodes) (Matousek, 1991; Nystrom et al., 1986). Collectively, these results indicate potential utility of beta oscillations as a sensitive marker of depression severity and duration.

### 1/f parameters exponent and offset are associated with depression severity

To our knowledge, we provide the first evidence of altered excitation-inhibition balance across a spectrum of depression severities via measurement with non-invasive EEG. Across datasets, we observed preliminary evidence of altered 1/f slope as a function of increased symptom severity at midline electrodes. The 1/f slope provides a marker of E/I balance, with flatter slopes (i.e., lower exponent values) reflecting a shift towards excitation, and steeper slopes (i.e., higher exponent values) a shift towards inhibition (Donoghue et al., 2020). Indeed, post-mortem tissue studies of individuals with depression indicate widespread changes in GABAergic and glutamatergic cell counts across regions/networks implicated in emotional regulation, such as prefrontal cortex and limbic regions (Hu et al., 2023). Our findings also align with previous longitudinal evidence demonstrating associations between symptom remission and 1/f slope, as measured via intracranial EEG in prefrontal and limbic regions (Hacker et al., 2025; Veerakumar et al., 2019). Collectively, our findings add to previous work indicating that altered excitation-inhibition balance underlies depression symptomology. We demonstrate the utility of the 1/f slope as a non-invasive measure of E-I balance to differentiate depression severities. Future work may expand on this finding to explore how the 1/f slope may be utilised as a physiological marker of treatment and recovery.

We also report an association between depression and the offset parameter of the 1/f slope, which indexes broadband shifts in power (Donoghue et al., 2020), and has been associated with neuronal spiking and fMRI measures of blood-oxygen-level-dependent activity (Manning et al., 2009; Winawer et al., 2013). It is plausible that our results reflect increases in excitability as a function of increased symptom severity. Interestingly, visual inspection of our data shows similar topographies between low gamma power and the 1/f offset, when correlated against depression symptoms. Indeed, a moderate correlation was observed between offset and low gamma power across electrodes. The strength of the association does suggest that either of these measures may provide a useful correlate of depressive symptomology, and raises the question of whether multiple EEG signatures are required to delineate depression severities. It will be important to determine the unique contribution of these measures to depression severity in future work.

### Limitations & future directions

This study has limitations which should be acknowledged. For instance, there were sampling differences across the two datasets analysed in this study. Participants in the TDBRAIN (2022) dataset were not required to have depression as their primary diagnosis, and some participants presented with other diagnoses such as obsessive-compulsive disorder and attention-deficit hyperactivity disorder, and/or recent history of medication use. However, depression is a mental health condition with a high degree of co-morbidity and a strength is that our findings relate to the population of individuals with a depression diagnosis. We observed similar trends across datasets (i.e., altered high-frequency oscillations) indicating that these EEG markers are robust to these sampling differences and may indeed be specific to depression symptomology. Further, we report a modest positive correlation between the 1/f slope offset and gamma power. It is methodologically challenging/impractical to remove the aperiodic component from high frequency oscillatory activity due to the absence of clear high-frequency peaks. As such, it is possible that these measures are inherently related and reflect a shared physiological process, rather than distinct aspects of brain function. Our results offer novel insights into the utility of non-invasive EEG markers of depression severity, but future work is required to delineate the extent to which gamma power and the spectral parameterisation model offer complementary markers of symptomology.

## Conclusion

In this study, we report evidence of altered high frequency oscillations and excitation-inhibition balance across a spectrum of depression severity. We show similar trends across two independent datasets, indicating that these non-invasive EEG measures may provide reliable markers of symptom progression. Future research should investigate how these markers shift in response to intervention, which may inform their utility in precision psychiatry approaches aimed and improved assessment and treatment.

## Acknowledgements

J.G.H is supported by a Research Training Program award from the Australian Department of Education. JC is supported by an Australian Research Council Future Fellowship (FT230100656). J.J.H is supported by an Australian Research Council Discovery Early Career Research Award (DE240101348). The funder played no role in study design, data collection, analysis and interpretation of data, or the writing of this manuscript. We acknowledge the efforts by the research teams that collected the data for the two datasets utilised in this study, and for making them freely available as open source resources.

## Author contributions

J.G.H, J.C., J.J.H designed and conducted data analysis for this study. J.G.H wrote the initial draft. All authors edited the manuscript. J.C and J.J.H supervised the research.

